# Cortical and thalamic afferent connectomes distinguish ACC subregions of the macaque brain

**DOI:** 10.64898/2026.03.03.709300

**Authors:** DC Myers, TM Love, JL Fudge

## Abstract

In human and nonhuman primates, the anterior cingulate cortex (ACC) is an interface between the “interoceptive” and “exteroceptive” domains. The ACC contains discrete subdivisions that are distinct in cytoarchitecture and connectivity, and also serve unique functional roles. The subgenual ACC (sgACC) is a key area for arousal state modulation and processing negative emotion. Importantly, the sgACC is dysregulated in major depression and a major target for neuromodulation therapies such as deep brain stimulation. In contrast, the perigenual ACC (pgACC) is important for a host of cognitive functions, including error monitoring and social decision-making. Thus, understanding the major sources of afferent input to the sgACC and pgACC is essential for elucidating functional modulation, including in major depression and other related disorders. We took a mesoscopic ‘connectomic’ approach to examine the balance of projections to sgACC and pgACC from two sources of glutamatergic input: the prefrontal cortex (PFC) and insula, and the thalamus (n=6 macaque monkeys). Using paired injections of retrograde tracers into the sgACC and pgACC and unbiased statistical clustering, we revealed that the ACC subdivisions are under the influence of vastly different cortical and thalamic ‘drivers’, i.e. heavily weighted inputs. We also found hierarchical rules (governed by cortical granularity and thalamo-cortical connectivity) by which PFC-ACC and thalamus-ACC circuits are organized. Overall, agranular PFC/insula cortices and associated thalamic nuclei are heavily weighted towards the sgACC, whereas pgACC receives a more even balance of afferents from agranular, dysgranular, and granular cortices and the thalamic nuclei with which they are associated.

**SIGNIFICANCE STATEMENT:** The ACC contains discrete subdivisions based on cytoarchitecture and connectivity, which serve unique functional roles. The sgACC and pgACC subdivisions receive many similar inputs based on neuroimaging work. Here, we leverage higher resolution retrograde tract-tracing in macaques to examine the relative weights and relationships of multiple cortical and thalamic afferents to each region. Using unbiased analyses of labeled cells, we conclude that the balance of afferent inputs shifts from a connectome dominated by agranular cortices and their thalamic partners in sgACC, to a more balanced afferent connectome in pgACC, represented by agranular, dysgranular, and granular cortices and their broader thalamic partners. These results facilitate interpretation of functional studies, and bridge understanding of the connectional basis of psychiatric disorders.

---

The anterior cingulate cortex (ACC) is implicated in a host of psychiatric disorders, including major depression (MDD). Patients with MDD exhibit altered ACC metabolic activity and functional connectivity. Therefore, treatments that regulate ACC activity have arisen as promising interventions for MDD, including deep brain stimulation (DBS) and transcranial magnetic stimulation (TMS) (Drevets et al., 1992; Mayberg et al., 1999; Mayberg et al., 2005; Liston et al., 2014; Batail et al., 2023; Duprat et al., 2025). Yet, the resolution of ACC circuits is incomplete, and depends on translational studies in nonhuman primates, the closest anatomic model. As in humans, the macaque ACC is highly complex, with subdivisions distinct in cytoarchitecture and function (Carmichael and Price, 1994; Vogt et al., 1995; Palomero-Gallagher et al., 2019). The subgenual ACC (sgACC), positioned ventral to the genu of the corpus callosum, comprises areas 25 and 14c (Carmichael and Price, 1994). These areas lack granular layer IV and have poorly differentiated layers II-III (Carmichael and Price, 1994). Functionally, the sgACC is implicated in processing negative emotion and sustaining emotional arousal (Rudebeck et al., 2014; Alexander et al., 2020; Amemori et al., 2024). The perigenual ACC (pgACC), comprising areas 32 and 24a/b, is rostro-dorsally adjacent to the sgACC. Areas 24a and 24b lack granular layer IV (Barbas and Pandya, 1989; Carmichael and Price, 1994). Area 32, positioned between areas 24b and 25, is composed of a rostral dysgranular region (32r) and a caudal agranular region (32c) (Carmichael and Price, 1994; Palomero-Gallagher et al., 2019). pgACC plays a greater role in cognitive functions than sgACC, including conflict monitoring and decision-making, particularly in social contexts (Amemori and Graybiel, 2012; Chang et al., 2013; Apps et al., 2016; Lockwood and Wittmann, 2018; Dal Monte et al., 2020; Simon IV and Rich, 2024).

Establishing maps of afferent control will advance understanding of ACC function and its role in psychiatric illnesses. The amygdala’s basal and accessory basal nuclei are key afferents of both sgACC and pgACC (Amaral and Price, 1984; Sharma et al., 2020). Beyond the amygdala, the ACC receives a host of other excitatory inputs. Tract-tracing studies have demonstrated that the prefrontal cortex (PFC) and insula send many afferent inputs to the ACC (Carmichael and Price, 1995, 1996; Ongur and Price, 2000; Heilbronner and Haber, 2014; Joyce and Barbas, 2018). The thalamus, massively enlarged in primates, is also a source of glutamatergic input. Known inputs to the ACC from individual tract-tracing studies include the mediodorsal and midline nuclei (Barbas et al., 1991; Carmichael and Price, 1996; Hsu and Price, 2007).

While valuable, examining circuits in isolation neglects the neural context in which brain circuits exist. A ‘connectomics’ approach acknowledges that brain region function depends on the strength of converging afferent systems seeking the same target (Cho et al., 2013; Yeo et al., 2013). This approach is especially relevant for understanding primate ACC circuitry due to its vast connections with the expansive cortical mantle. Therefore, we took a ‘meso-connectomics’ view to examine the sgACC and pgACC, quantifying retrogradely labeled neurons throughout the PFC, insula, and thalamus, then locating their position in cortex (Cho et al., 2013; McHale et al., 2022) and in thalamic nuclear zones (Jones, 1998; Garcia-Cabezas et al., 2023). An unbiased clustering approach was used to examine the main cortical and thalamic ‘drivers’ of ACC subdivisions. Individual afferent inputs to sgACC and pgACC were then examined within and across animals using correlative analyses to determine how specific combinations of inputs project across the ACC trajectory. Finally, we compared all afferents to each region using cortical laminar differentiation criteria, and assessed all thalamic inputs based on their known connections with the cortex. The afferent ‘connectome’ to each region followed specific rules related to cortical granularity and thalamo-cortical projections. Thus, while sgACC and pgACC receive many similar inputs, the relative weighting of the cortical and thalamic connectome shifts predictably between them, and is governed small differences in sgACC and pgACC laminar organization.

## Materials and Methods

### Study Design

6 male *Macaca fascicularis* (weight range = 4.2 – 5.5 kg, age range = 4 – 5 years) were used in this study (Worldwide Primates, Tallahassee, FL, USA). Some animals used in a previous study (Sharma et al., 2020). In each animal, paired injections of different retrograde neuronal tracers were placed into sgACC (BA 25/14c) and pgACC (BA 32/24a,b) of the same hemisphere using stereotaxic techniques. After brain processing, retrogradely labeled neurons were charted and quantified throughout the entire extent of the PFC/insula and thalamus, and then localized into specific cortical and thalamic regions using adjacent sections stained for relevant markers. The retrogradely labeled neuron counts for each afferent site were normalized to total labeled cells in either the PFC and insula, or thalamus, for each injection in each animal. Using these proportions permitted assessment of the relative weight of specific afferent paths to sgACC and pgACC in each case.

### Surgical Procedures and Tissue Preparation

All surgeries conducted for this study were approved by the University of Rochester Medical Center Committee on Animal Research (UCAR) and followed National Institute of Health guidelines. To determine surgical coordinates for tracer injection into sgACC and pgACC, each animal received a T2 anatomic MRI prior to surgery. MRI images were used to identify the position of areas the sgACC and pgACC on coronal sections spaced 0.9 mm apart in each individual animal. Therefore, each animal had unique coordinates for sgACC and pgACC injections. During the 3 days preceding stereotaxic surgery, animals received a daily oral dose of gabapentin (25 mg/kg) for pain control, which was maintained for 3 days post-surgery. Prior to stereotaxic surgery, animals received an intramuscular injection of ketamine (10 mg/kg), were intubated, and maintained on isoflurane gas during the procedure. Animals were stabilized in a stereotaxic frame and a craniotomy calculated to accommodate all injections was performed under sterile conditions. In all animals, a small hole was made in the dura and injections (40 nL) of different retrograde neuronal tracers (either fluoro-Ruby [FR], fluorescein [FS], or lucifer yellow [LY]) were pressure-injected (sterilized Hamilton syringe, 0.5 μL) into the brain at pre-determined coordinates over a 10-minute period into sgACC and pgACC of the same hemisphere. Each tracer type was only used for one injection per animal. After each injection, the syringe was left in place for 20 minutes to prevent backflow of tracer up the injection track. After all injections were carried out, the bone flap was replaced and the surgical site was sutured.

Animals were maintained on standard post-operative observation protocol. Two weeks post-surgery, animals were deeply anesthetized with pentobarbital (20 mg/kg) and euthanized via intracardiac perfusion. Perfusions were performed using 0.9% saline containing 0.5 mL of heparin sulfate (200 mL/minute for 15-20 minutes), which was immediately followed by cold 4% paraformaldehyde in 0.1M PB (200 mL/minute for a total of 6L). Brains were extracted and fixed overnight in 4% paraformaldehyde solution, and then were sequentially submerged in 10%, 20%, and 30% sucrose solutions until they sank in each. Then, 40 μm sections through the entire brain were collected using a freezing, sliding microtome. Sections were stored in serial compartments in cryoprotectant solution and stored at −20°C.

### Histology

#### Immunocytochemistry (ICC)

To identify injection sites and retrogradely labeled neurons throughout the PFC, insula, and thalamus, at least 1:12 near adjacent sections through the cortex and thalamus were immunostained for each tracer type. Anti-sera for tracers are not cross-reactive based on previous studies (Haber et al., 2000; Sharma et al., 2020; McHale et al., 2022). Selected sections were first rinsed in phosphate buffer with 0.3% Triton-X (PB-TX) overnight. The next day, sections were treated with an endogenous peroxidase inhibitor for 5 minutes and rinsed again in PB-TX for 6 x 15 minutes. Then, sections were pre-incubated in a blocking solution of 10% normal goat serum in PB-TX for 30 minutes. Following this, sections were incubated in primary antisera to FR (1:1000, Invitrogen, Carlsbad, CA, rabbit), FS (1:2000, Invitrogen, rabbit), or LY (1:2000, Invitrogen, rabbit) at 4°C for 4 nights on a rocker. Sections were then rinsed thoroughly (6 x 15 minutes) in PB-TX. Following this, sections were blocked again with 10% normal goat serum in PB-TX and incubated for 40 minutes in biotinylated secondary anti-rabbit antibody diluted in 10% normal goat serum (1:200, Vector Laboratories, Newark, CA). After incubation in secondary antibody for 40 minutes, sections were thoroughly rinsed in PB-TX 6 x15 minutes, then incubated in an avidin-biotin complex (Vectastain ABC Kit, Vector Laboratories) for 60 minutes. After rinses in PB, sections were visualized with 3,3-diaminobenzidine (DAB, 0.05 mg/mL in 0.1 M Tris buffer). Sections were then rinsed overnight and mounted out of a mounting solution (0.5% gelatin in 20% EtOH in double distilled water) onto gelatin-coated slides. After mounting, sections were dried for 3 days, and then coverslipped with DPX mounting media (Electron Microscope Sciences, Hatfield, PA).

#### Cresyl Violet

To identify cytoarchitectonic boundaries of cortical areas, sections that were adjacent with tracer-labeled sections, were stained with cresyl violet solution. Briefly, sections were rinsed in PB, mounted onto gelatin-coated slides and dried on a warmer. Then slides were dehydrated through increasing gradients of alcohol and cleared in xylenes for 48 hours. Slides were subsequently rehydrated though decreasing alcohol baths and air-dried until ready for cresyl violet processing. Once stained in cresyl violet (Chroma-Gesellschaft, Stuttgart, Germany), slides were dried on a warming plate and coverslipped.

#### Acetylcholinesterase (AChE)

To identify thalamic nuclear boundaries, additional adjacent sections throughout the thalamus for each case were stained for AChE (Cavada et al., 1995). Sections were chosen, rinsed thoroughly in PB, and stained for 2-2.5 hours using the Geneser-Jensen method (Geneser-Jensen and Blackstad, 1971).

### Analysis

#### Injection Site Placement

The center of the injection site within the sgACC or pgACC was drawn under a microscope at low power on at least 1:12 sections using Neurolucida software. Adjacent cresyl violent-stained sections were used to determine the center of the injection site placement within the sgACC and pgACC based on cytoarchitectonic profiles. Cases with leakage into white matter or outside the ACC were excluded from the analysis. A whole brain survey of known connections was also used to confirm injections site placement. For example, area 25 projects to the ventromedial shell striatum, and area 32 does not (Haber et al., 2006). In contrast, area 32 projects to the central and lateral sections of the ventral striatum (Choi et al., 2017). We leveraged both anterograde and retrograde properties of the tracers to confirm injection site placement, although the retrograde labeling in the cortex and thalamus were used for this study.

#### Charting of Retrogradely Labeled Cells in Cortical and Thalamic Regions

Retrogradely labeled cells in the PFC, insula, and thalamus were manually charted under brightfield microscopy on a 1:24 series of slides (Olympus AX70 microscope, Olympus, Tokyo, Japan) interfaced with Neurolucida 360 software (Microbrightfield Bioscience, Williston, VT). Retrogradely labeled cells were charted throughout the rostral-caudal extent of the PFC and insula (12-14 sections), and thalamus (11-12 sections), along with landmarks (blood vessels, white matter tracts, etc.) to permit alignment with adjacent sections stained with cresyl violet and AChE. These were also charted using a 10x objective under bright-field illumination, and included fiducial markers including fiber tracts and blood vessels. Charts of retrogradely labeled cells and adjacent cresyl violet or AChE-stained sections were then saved as vector graphics, imported to Adobe Illustrator, and aligned in transparent layers using traced landmarks. This enabled the creation of laminar maps and templates of thalamic regions for each case.

#### Localization of retrogradely labeled neurons

Cortical and insula regions containing retrogradely labeled cells were divided into cytoarchitectonic regions using the criteria of Carmichael and Price (Carmichael and Price, 1994) **(Fig. 1A**), and retrogradely labeled cells in the thalamus were first localized in individual thalamic nuclei **(Fig. 1B**), which were then assigned to canonical thalamic nuclear groups based on similar connectivity and function (Goldman-Rakic and Porrino, 1985; Jones, 1985; Ilinsky and Kultas-Ilinsky, 1987; Hsu and Price, 2007; Garcia-Cabezas et al., 2023)**(Fig. 1C).** The number of cells identified in each cortical and thalamic area (e.g. **Fig. 1D**) was then quantified for each injection site and assembled on spreadsheets. Since the cortex and thalamus are distinct brain regions (i.e. one laminar, one nuclear) with different dispersions and packing of cells, the percentage of labeled cells for cortical areas and thalamic regions for each injection site was calculated separately as a percentage of labeled cells in each structure. While all sections were counted, we eliminated sections containing the center of the injection site in the analysis of cortical cell counts to conservatively exclude labeled cells at the injection site.

**Figure 1.**
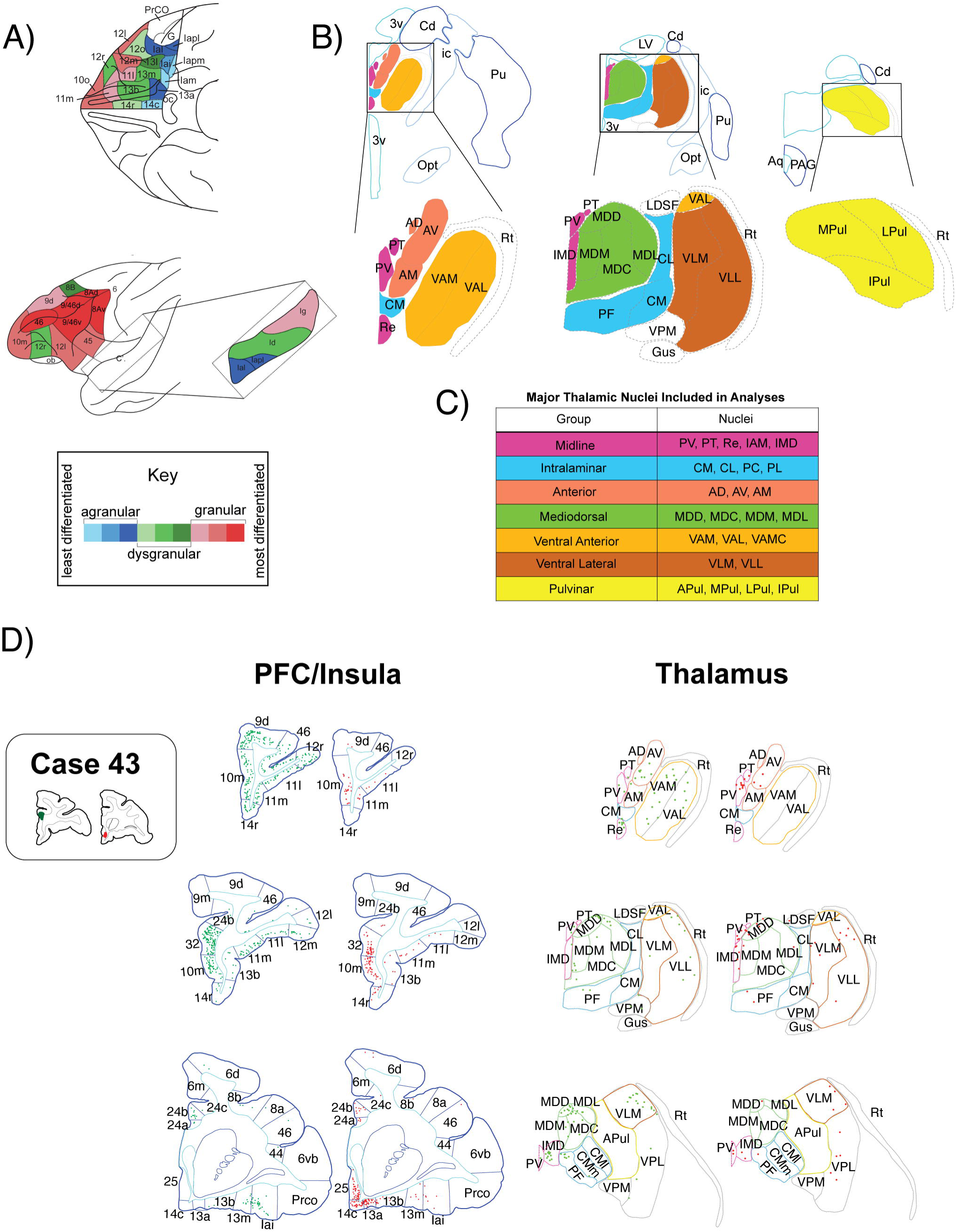
Charting retrogradely labeled cells in the PFC, insula, and thalamus. A. Schematic of cortical subregions on the orbital and lateral surface of the cortex based on Carmichael and Price, 1994. Color coded for laminar differentiation according to McHale et al, 2022. Inset: insula of the Sylvian fissure. B. Schematic of thalamic nuclei from rostral to caudal (left to right). C. Thalamic nuclear regions based on cortical connections. D. Representative charts of retrogradely labeled cells after dual injections in Case 43 (red, sgACC; green, pgACC). Not all charts are shown. *PV = paraventricular nucleus, PT = paratenial nucleus, Re = reuniens, IAM = interanteromedial nucleus, IMD = intermediodorsal nucleus, CM = centromedian nucleus, CL = centrolateral nucleus, PC = paracentral nucleus, PF = parafascicular nucleus, MDD = dorsal mediodorsal nucleus, MDC = central mediodorsal nucleus, MDM = medial mediodorsal nucleus, MDL = lateral mediodorsal nucleus, VAM = medial ventral anterior nucleus, VAL = lateral ventral anterior nucleus, VAMC = magnocellular ventral anterior nucleus, VLM = medial ventral lateral nucleus, VLL = lateral ventral lateral nucleus, APul = anterior pulvinar, MPul = medial pulvinar, LPul = lateral pulvinar, IPul = inferior pulvinar*.

#### Cluster Analysis of Normalized Cell Count Data

To identify groups of cortical and thalamic areas with similarly weighted projections across ACC injections, k-means cluster analyses were done on normalized cell count data. K-means cluster analysis is an unbiased machine learning algorithm that partitions data into distinct groups such that similar data points cluster together and dissimilar data points are clustered into separate groups (Kaufman and Rousseeuw, 2009). Three cluster plots were constructed: 1 for all 12 injection sites (ACC cluster), 1 including data from sgACC injection sites only (sgACC cluster), and 1 including data from pgACC injection sites only (pgACC cluster).

Cell counts for cortical and thalamic areas were z-scored using the formula (x - μ) / σ, where x is the number of cells identified in a cortical or thalamic area for one case, μ is the mean cell count for all cortical or thalamic areas in that case, and σ is the cell count standard deviation for all cortical or thalamic areas in that case. Cortical cell count data were z-scored based on cortical means and standard deviations only, and thalamic cell count data were z scored based on thalamic means and standard deviations only. This approach ensured that cluster plots were not influenced by variation in cell count totals across injection sites or between cortical and thalamic cell counts.

To ensure that the data were not being underclustered or overclustered, we used the silhouette method to determine the optimal number of clusters for our dataset (Rousseeuw, 1987). The silhouette method determines the average silhouette width for increasing numbers of clusters, where a higher silhouette width indicates higher intragroup similarity and lower intergroup similarity. We chose the number of the clusters with the highest average silhouette width. Cluster and silhouette analysis were conducted in RStudio using the “factoextra” and “cluster” libraries. All code is publicly available at https://doi.org/10.6084/m9.figshare.31395261. The patterns of z-scores across injection sites for cortical and thalamic areas were examined in order to properly interpret each cluster. Then, descriptive names for the clusters were created according to their interpretation. Clusters were then reported in anatomic space by color-coding PFC and thalamus subregions involved in each cluster.

#### Variable Projections to Injection Sites in ACC

We made spider plots (RStudio, “fmsb” and “scales” libraries) of the percentage of labeled cells for each injection in each case to represent their relative distribution across main cortical and thalamic subregions. Because the goal was to examine variability across injection sites, each injection site within sgACC or pgACC was represented on the same spider plot for both cortex and thalamus.

#### Common Inputs to sgACC and pgACC Across Injection Sites

As another method for extracting the major sources of input across ACC subregions, we used the Tukey-Fence method. This permitted identification of cortical and thalamic areas that were large ‘outliers’ for sgACC sites or pgACC sites (compared to all sites in the group), i.e. had abnormally large projections compared to other cortical and thalamic areas. First, we used z-scored cell count data to calculate Euclidian distance for all cortical and thalamic areas for sgACC and pgACC separately, using the formula Distance 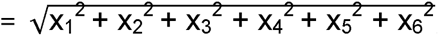, where x represents a cortical/thalamic area’s z-score for each injection site. After calculating Euclidian distance for all cortical and thalamic areas for both sgACC and pgACC, we determined the interquartile range (IQR) and the first (Q1) and third (Q3) quartiles for sgACC and pgACC. Then, all Euclidian distances for both sgACC and pgACC were compared to an “upper fence” that was calculated with the formula upper fence = Q3+ 1.5 × IQR, which represents a threshold for determining high statistical outliers. Areas that were and were not high statistical outliers for sgACC and pgACC were represented with a bar plot.

#### Analysis of Networks Across a Linearized Axis of the ACC

We aimed to evaluate whether cortical or thalamic areas project in a systematic pattern across ACC areas 25, 32, and 24. To do this, we first identified the location of each injection site in the ACC on a standardized macaque brain atlas for 4-5 year old animals, as done previously (Sharma et al., 2020). The ACC wraps around the genu of the corpus callosum in primate species due to general expansion and folding of the frontal lobe (van Heukelum et al., 2020). Therefore, we ‘unfurled’ the ACC so that area 32r, which is the most rostral area along the ACC curvature formed a pivot-point, and areas dorsal-caudal to this were rotated 180 degrees. This created a linearized ACC where area 24 was positioned at one end, followed by area 32r, then 32c, and finally by areas 25 and 14c. This allowed us to examine how projections shift across subdivisions of the ACC in a linear manner, closer to the organization in lower species. We used linear regression to quantify the relationship between percentage of total inputs (y axis) and position of the injection site on a flattened ACC (x axis) for all cortical and thalamic areas.

#### Correlated Cortical-Cortical and Cortical-Thalamic Afferents Systems

To determine patterns in cortical-cortical, thalamic-thalamic, or cortico-thalamic co-projections to the ACC, Spearman correlations were performed for the normalized cell counts between regions. Cortical and thalamic areas were selected for comparison to other cortical and/or thalamic areas based on inspection of similar relative contributions to the ACC (spider plots).

#### Weighting of Diverse Cortical and Thalamic Afferents (’Connectomes’) to sgACC versus the pgACC

Finally, we used chord plots to examine the relative contribution of diverse cortical and thalamic areas to the sgACC versus pgACC across animals. Separate chord plots of all weighted inputs were made for cortical and thalamic projections. Cortical areas were considered either “agranular”, “dysgranular”, or “granular” based on the classifications described in previous work, and were color coded as such. This was done to visualize how increments of cortical granularity might shape projection paths to sgACC and pgACC as it does in other downstream structures (Cho et al., 2013; McHale et al., 2022). On the basis of the known reciprocal connectivity of thalamic nuclei with the PFC (Goldman-Rakic and Porrino, 1985; Jones, 1985; McFarland and Haber, 2002; Hsu and Price, 2007), thalamic nuclear groups were color-coded with respect to their main connectivity with agranular, dysgranular, and granular regions of cortex. All chord plots were made in RStudio using the “circlize” library. Cortical and thalamic areas were represented as a node on the chord plot, with tick marks showing the average percentage of inputs from each area to sgACC and pgACC.

## Results

### Injection site placement in the ACC

Injection pairs along the ACC cytoarchitectural gradient (**Fig. 2A**) are depicted in **Fig. 2B**. A summary of the tracers used and the hemispheres targeted in each case is provided in **Fig. 2C**, color-coded for pairs in the same animal. Five out of six injections into the sgACC included area 25, with one injection (Case 53FR orange) straddling areas 25 and 14c. One injection (Case 39FR, pink) was in area 14c alone. In pgACC, injections were at various levels of area 32 (Cases 49FR, green, Case 43LY, red, and Case 53FS, red). In Cases 49FR and 43LY, the injection sites were in area 32c. In Case 53FS, the injection site straddled areas 32r and 32c. Three injections were in area 24 (Cases 39LY, pink, 46FR blue, and 50FS, yellow). Cases 39LY and 46FR were entirely in area 24b, and Case 50FS was in area 24a/b. Representative photomicrographs of injection sites for Case 43 (areas 25 and 32r), and Case 50 (areas 25 and 24b) are shown (**Fig. 2D**).

**Figure 2.**
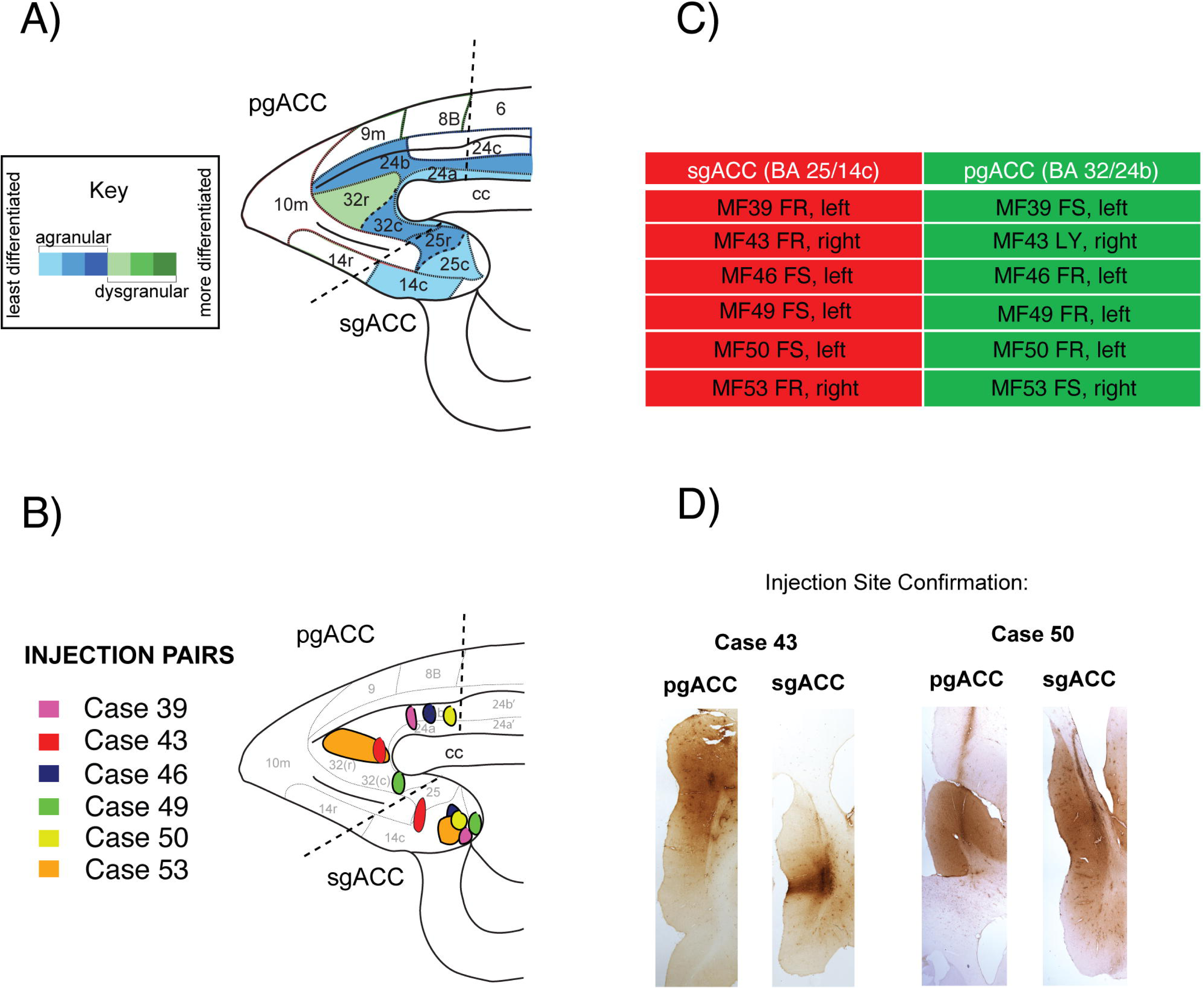
Summary of injection pairs. A. Sagittal view of the ACC, color coded by granularity assignment (from McHale et al., 2022) with sgACC and pgACC boundariesB. Sagittal view the ACC, showing injection pairs. C. Summary of injection pairs. D. Low power images of racer-stained coronal sections through the injection sites in Case 43 and Case 50. Dark lines in Case 50 images reflects DAB-stained gliosis.

### Unbiased clustering analysis of cortical and thalamic projections to ACC

We first examined data from all 12 injection sites combined, without regard to placement in ACC subregions. Silhouette analysis revealed that 3 clusters was best for data separation (**Fig. 3A**). In the ACC analysis, one cluster was composed of areas with large z-scores for several injection sites, including cortical areas 25, 14c, 13a, 13b, 32c, and 10m, the midline and intralaminar thalamic nuclei (**Fig. 3B, red**). A second, cluster composed of cortical areas 24b, 32r, 9, and 46, and the mediodorsal (MD) and ventral anterior (VA) thalamic nuclei, also had large z-scores for several injection sites (**Fig. 3B, blue**). The remaining cortical and thalamic cluster were characterized by z-scores near or below mean for all injection sites (**Fig. 3B, green**). We named the red cluster ‘arousal’ due to inclusion of structures required for alerting and interoceptive function (area 25, midline thalamus). We named the blue cluster ‘executive’ due to inclusion of classic networks involved in working memory, and the green cluster was labeled ‘modulatory’ due to the inclusion of regions that made relatively small contributions to the ACC as a whole.

**Figure 3.**
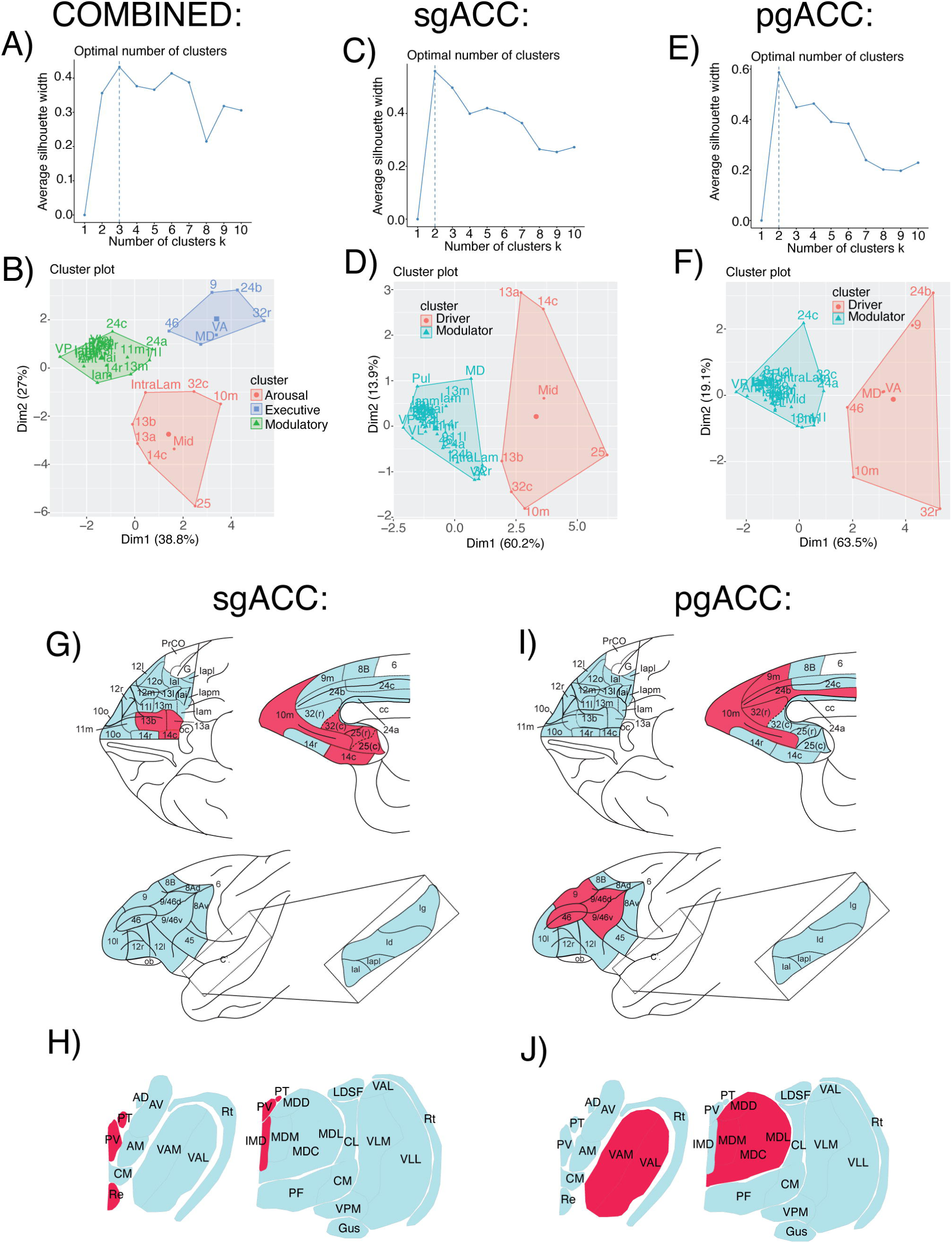
Unbiased clustering of cortical and thalamic afferents. A. Silhouette analysis for all ACC injection sites. B. Cluster plot for ACC injection sites, revealing 3 clusters which were interpreted as ‘arousal’, ‘executive’ and ‘modulatory’ inputs (see text). C. Silhouette analysis for sgACC sites. D. Cluster plot for sgACC sites, revealing 2 main clusters interpreted as ‘driver’ and ‘modulator’ afferents. E. Silhouette analysis for pgACC sitesF. Cluster plot for pgACC sites, revealing 2 main clusters also termed ‘driver’ and ‘modulator’. G-J. Anatomic maps depicting ‘driver’ (red) and ‘modulator’ (teal) sites for sgACC (G, H) and pgACC (I, J).

K-means clustering for sgACC sites alone **(Fig. 3C-D**) versus pgACC sites (**Fig. 3E-F**) each revealed clear 2-cluster peaks on silhouette analyses (**Fig. 3C, E**). For the sgACC sites, one cluster was composed of areas with large z-scores for sgACC sites (red), and another cluster was composed of areas with near or below mean z-scores for sgACC sites (teal)(**Fig. 3D**). The red cluster therefore was interpreted to contain the most heavily weighted prefrontal and thalamic inputs of the sgACC: areas 25, 14c, 13a, 13b, 32c, 10m, and midline thalamic nuclei **(Fig. 3D**). We refer to these as “drivers”. The teal cluster, containing regions with z-scores at or below mean, were labeled “modulators” of sgACC. Likewise, for pgACC, we elucidated one cluster composed of “drivers” based on their large z-scores for pgACC sites. These were: areas 24b, 32r, 10, 9, 46, and the MD and VA thalamus (**Fig. 3F**). All remaining prefrontal cortical and thalamic areas projecting to the pgACC clustered together based on their near or below mean z-scores, and were considered “modulators” of pgACC (**Fig. 3F**). It is notable that area 32c is a “driver” of sgACC (Fig. 3D), but areas 25/14c are not “drivers” of the pgACC (Fig. 3F), suggesting that, at least for ‘drivers’, the intrinsic flow of information within the ACC is more “top-down” than “bottom-up”.

Mapping the individual constituents of driver and modulators in anatomic space with drivers demarcated in red and modulators demarcated in teal revealed largely different cortical and thalamic drivers for sgACC and pgACC sites (**Fig. 3G-J**). On the orbital surface, areas 13a, 13b, and 14c were drivers of sgACC, whereas no areas were drivers of the pgACC (**Fig. 3G, H**). On the medial wall, sgACC drivers predominate along the ventromedial cortex while pgACC drivers occupy the dorsomedial cortical regions. Area 10m was a driver of both sgACC and pgACC. In the dorsolateral cortex, areas 9 and 46 were drivers of pgACC, with no regions forming a driver relationship in the sgACC. Surprisingly, the insula had only modulatory level inputs to both the sgACC and pgACC. Thalamic nuclei subregion drivers to each region were clearly separated, with the midline thalamic nuclei providing strong inputs to the sgACC sites, and MD and VA nuclei forming to the strongest inputs to pgACC sites (**Fig. 3I, J**).

### Among key sgACC and pgACC drivers lies variability

Variation in injection site location within the sgACC and pgACC resulted in variation in the distribution of retrogradely labeled cells in some PFC and insula subregions (**Fig. 4A**, **Table 1**). For the sgACC sites, areas 25, 14c, and 13a/b consistently formed a large proportion of inputs across sgACC sites, but shifts in the proportions of other inputs were found. For example, in cases with injections restricted to area 25 (Cases 43FR, 50FS, 46FS, and 49FS), area 10 was a prominent input, whereas an injection restricted to area 14c (Case 39FR), resulted in a lower proportion of labeled cells in area 10, and relatively more labeled cells in area 13 (over 35% of all cortical projections). Case 53FR (including both area 25 and 14c) also had lower proportions of labeled cells in area 10, and received more inputs from area 24 compared to other sgACC sites.

**Figure 4.**
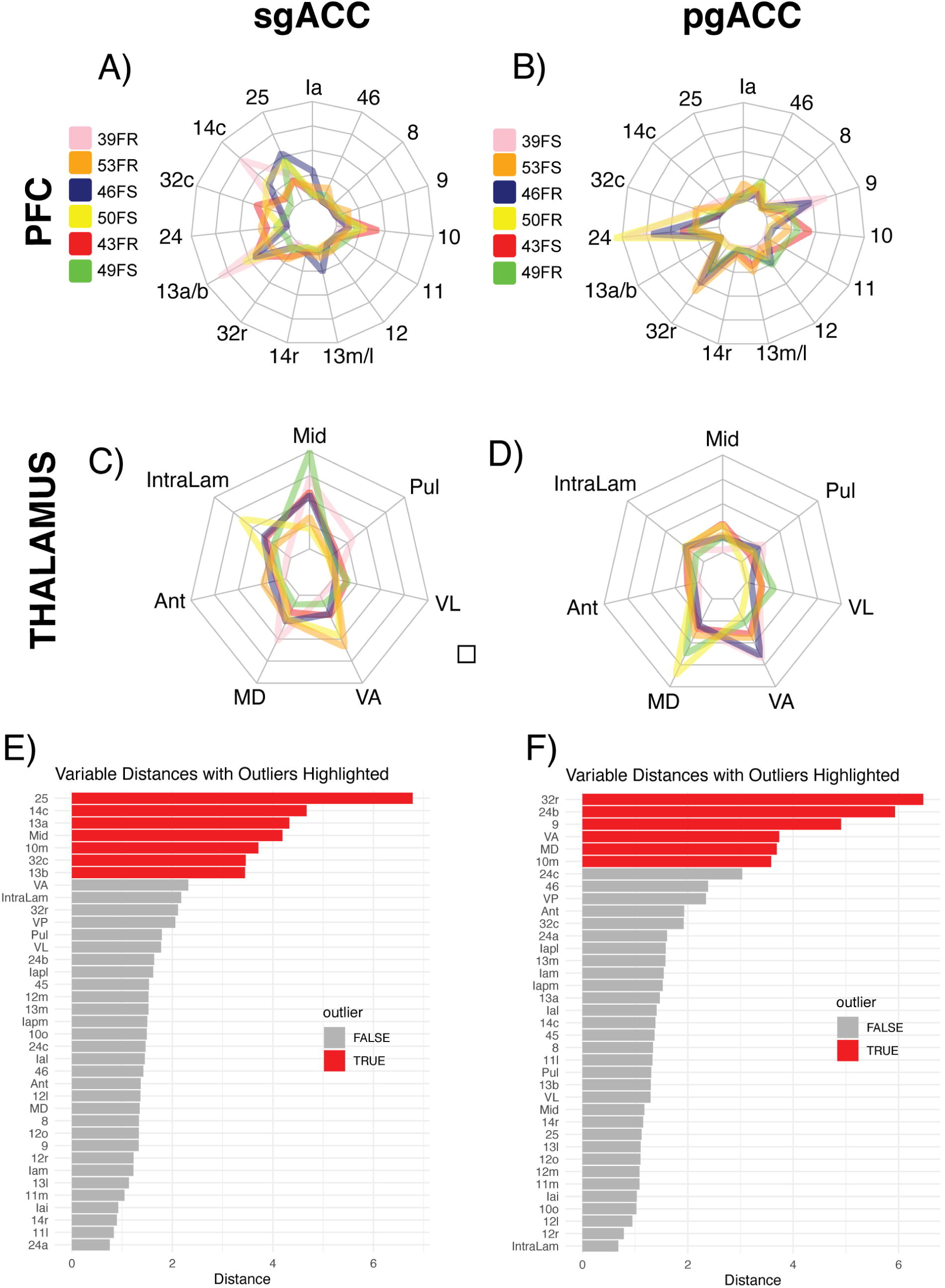
Variability of afferent networks within sgACC and pgACC. A-D. Spider plots showing the distribution of retrogradely labeled cells throughout the PFC after injection into sgACC (A, B) and pgACC (C, D). Each polygon represents one injection site. Scale max = 60% of all labeled PFC cells, Scale max = 45% of all labeled thalamus cells. E-F. Bar plots showing PFC and thalamic areas identified as high statistical outliers for sgACC sites (E) and pgACC sites (F).

**Table 1.** Normalized numbers of labeled cells in cortical (top) and thalamic (bottom) regions for each injection site.

| **sgACC** | **la** | **25** | **14c** | **32c** | **24** | **13a/b** | **32r** | **14r** | **13m/l** | **12** | **11** | **10** | **9** | **8** | **46** |
| --- | --- | --- | --- | --- | --- | --- | --- | --- | --- | --- | --- | --- | --- | --- | --- |
| **Case 43FR - Area 25** | 2.66 | 10.58 | 5.68 | 15.72 | 9.92 | 19.29 | 8.10 | 2.24 | 2.18 | 0.24 | 2.42 | 18.80 | 0.54 | 0.06 | 0.18 |
| **Case 46FS - Area 25** | 13.01 | 24.11 | 15.73 | 3.54 | 0.60 | 19.32 | 2.44 | 2.54 | 11.33 | 0.45 | 1.00 | 5.89 | 0.01 | 0.01 | 0.02 |
| **Case 49FS - Area 25** | 1.62 | 20.99 | 2.37 | 1.39 | 5.78 | 2.08 | 4.22 | 0.00 | 2.37 | 3.82 | 5.49 | 10.53 | 2.66 | 1.74 | 3.18 |
| **Case 50FS - Area 25** | 3.16 | 19.99 | 17.38 | 11.50 | 3.57 | 24.66 | 2.46 | 0.55 | 0.65 | 0.15 | 0.95 | 13.06 | 0.85 | 0.40 | 0.35 |
| **Case 53FR - Area 25/14c** | 5.52 | 9.88 | 6.75 | 10.58 | 12.87 | 10.85 | 9.43 | 2.24 | 4.81 | 4.21 | 2.94 | 5.46 | 6.56 | 0.84 | 6.59 |
| **Case 39FR - Area 14c** | 4.24 | 13.30 | 32.14 | 3.62 | 1.19 | 37.58 | 0.06 | 1.53 | 1.19 | 0.40 | 0.96 | 3.06 | 0.68 | 0.00 | 0.00 |
| **pgACC** |  |  |  |  |  |  |  |  |  |  |  |  |  |  |  |
| **Case 39FS - Area 24a/b** | 0.70 | 1.74 | 4.54 | 10.19 | 42.71 | 0.42 | 0.63 | 0.07 | 0.07 | 2.02 | 0.70 | 0.42 | 28.40 | 2.51 | 3.56 |
| **Case 46FR - Area 24a/b** | 0.47 | 0.00 | 0.16 | 1.26 | 30.65 | 0.95 | 22.20 | 0.79 | 1.50 | 10.58 | 1.66 | 2.84 | 20.85 | 0.79 | 4.58 |
| **Case 50FR - Area 24a/b** | 0.00 | 0.13 | 0.08 | 8.55 | 47.91 | 1.18 | 1.14 | 0.00 | 1.23 | 7.66 | 2.20 | 0.93 | 14.30 | 2.16 | 9.61 |
| **Case 53FS - Area 32r/c** | 7.77 | 0.48 | 1.60 | 11.72 | 10.94 | 1.11 | 27.05 | 3.30 | 10.83 | 2.28 | 4.86 | 9.94 | 3.24 | 0.00 | 4.88 |
| **Case 43LY - Area 32r** | 2.88 | 1.11 | 0.35 | 1.91 | 17.17 | 0.80 | 14.64 | 1.95 | 8.16 | 5.06 | 9.32 | 19.57 | 8.07 | 0.31 | 8.39 |
| **Case 49FR - Area 32c** | 1.83 | 2.78 | 0.05 | 3.92 | 13.76 | 0.73 | 15.55 | 0.39 | 2.94 | 11.49 | 9.50 | 14.17 | 11.28 | 0.05 | 10.60 |

| **sgACC** | **Midline** | **Intralaminar** | **Anterior** | **Mediodorsal** | **VA** | **VL** | **Pulvinar** |
| --- | --- | --- | --- | --- | --- | --- | --- |
| **Case 43FR - Area 25** | 34.93 | 19.65 | 5.24 | 11.79 | 13.97 | 8.30 | 6.11 |
| **Case 46FS - Area 25** | 32.90 | 21.07 | 8.92 | 18.12 | 12.75 | 1.60 | 4.65 |
| **Case 49FS - Area 25** | 59.38 | 13.28 | 3.52 | 6.64 | 5.86 | 9.38 | 1.95 |
| **Case 50FS - Area 25** | 12.66 | 38.61 | 3.16 | 15.82 | 27.85 | 1.90 | 0.00 |
| **Case 53FR - Area 25/14c** | 19.39 | 11.91 | 14.27 | 17.17 | 35.32 | 1.25 | 0.69 |
| **Case 39FR - Area 14c** | 43.08 | 2.56 | 1.54 | 31.28 | 1.03 | 1.54 | 18.97 |
| **pgACC** |  |  |  |  |  |  |  |
| **Case 39FS - Area 24a/b** | 1.65 | 12.24 | 0.00 | 20.24 | 40.47 | 8.00 | 17.41 |
| **Case 46FR - Area 24a/b** | 9.56 | 14.69 | 3.28 | 18.26 | 38.52 | 3.14 | 12.55 |
| **Case 50FR - Area 24a/b** | 17.06 | 14.90 | 3.73 | 51.76 | 11.37 | 0.98 | 0.20 |
| **Case 53FS - Area 32r/c** | 11.04 | 12.03 | 6.40 | 24.89 | 26.88 | 9.55 | 9.20 |
| **Case 43LY - Area 32r** | 17.81 | 13.97 | 7.12 | 20.27 | 24.11 | 10.14 | 6.58 |
| **Case 49FR - Area 32c** | 9.74 | 6.15 | 3.59 | 37.44 | 15.90 | 17.44 | 9.74 |

For pgACC injection sites (**Fig. 4B**, **Table 1**), area 24a/b were consistently large inputs. However, areas 10, 12 and 32r inputs were somewhat more variable across injection sites. Higher proportions of labeled cells in area 9 were associated with injections in area 24a/b, with lesser proportions associated with area 32 injection sites (Cases 49FR, 53FS, and 43LY).

All sgACC injections had relatively high proportions of labeled cells in the midline nuclei, with more variable proportions of labeled cells in other nuclear regions **(Fig. 4C**, **Table 1**). For example, in Case 39FR, where the injection site was confined to area 14c, high proportions of labeled neurons in the midline thalamic nuclei were complemented by high proportions of labeled cells in the MD thalamus and the pulvinar. In contrast, injections restricted to area 25 (Cases 43FR, 46FS, 49FS, and 50FS) had relatively fewer inputs from these nuclear regions. Instead, relatively higher proportions of labeled neurons were found in the intralaminar nuclei for these cases. Case 53FR, which straddled areas 25 and 14c had more evenly distributed percentages of labeled neurons across these thalamic regions, consistent with a broader array of inputs. In the pgACC sites **(Fig. 4D**, **Table 1**), all injection sites had relatively high levels of labeled neurons in the MD and VA thalamus, with relatively less involvement of the midline thalamus. Area 24a/b injections in Cases 39FS and 46FR resulted in higher proportions of labeled cells in the VA thalamus (38.5-40.4% of all thalamic inputs to pgACC) than area 32 injections which had relatively even proportions of labeled neurons in MD and VA.

To quantify the main projections to each region amidst variability, we segregated high statistical ‘outliers’ of the PFC and thalamus inputs for sgACC and pgACC sites using a Tukey’s fence analysis. High statistical ‘outliers’ represented areas with large projection weights across injection sites, and for sgACC were areas 25, 14c, 13a/b, 10m, 32c, and the midline thalamic nuclei (**Fig. 4E, red**). High statistical ‘outliers’ for pgACC were areas 32r, 24b, 9, and MD and VA thalamus (**Fig. 4F, red**). This analysis generally confirms ‘driver’ results (heavily weighted inputs) from unbiased clustering. However, it is important to understand that individual variability in injection site placement, and shifts in relative contributions of individual cortical and thalamic regions are present. In this regard, sgACC injection sites appeared to have higher variability in afferent inputs compared to pgACC sites **(Fig. 4A, B).**

### Cortical and thalamic projection correlations along the ACC trajectory

To further understand injection site variability noted above in terms of the cytoarchitectural trajectory of the ACC, we assessed the relative placement of injection sites from the caudal sgACC (most undifferentiated) to the dorsal pgACC (most differentiated) sites. To do this, we placed injections sites from blocked and immunostained sections in a standardized A-P space for 4 year old animals (∼4kg) (Sharma et al., 2020), then we ‘cut’ and ‘unfolded’ the ACC map, converting standardized A-P values into values on a scale where 0 is the most caudal injection site in the sgACC (case 49FS) (**Fig. 5A**). Using these values, we found PFC and thalamic areas with proportions of labeled cells that increased or decreased incrementally along the ACC. Linear decreases from 3 PFC and one thalamic region were seen moving from the sgACC to the pgACC (**Fig. 5 B-F**). These areas were: area 25 (R^2^ = 0.7153, p = 0.000323), area 14c (R^2^ = 0.2835, p = 0.0432), area 13a/b (R^2^ = 0.3322, p = 0.0291), and the midline thalamic nuclei (R^2^ = 0.4501, p = 0.0101). In contrast, 2 PFC areas primarily exhibited a linear increase in the weight of their projections when moving from the sgACC to the pgACC. These areas were: area 24a/b (R^2^ = 0.7825, p < 0.0001), and area 9 (R^2^ = 0.5472, p = 0.0036)**(Fig. 5G-H)**.

**Figure 5.**
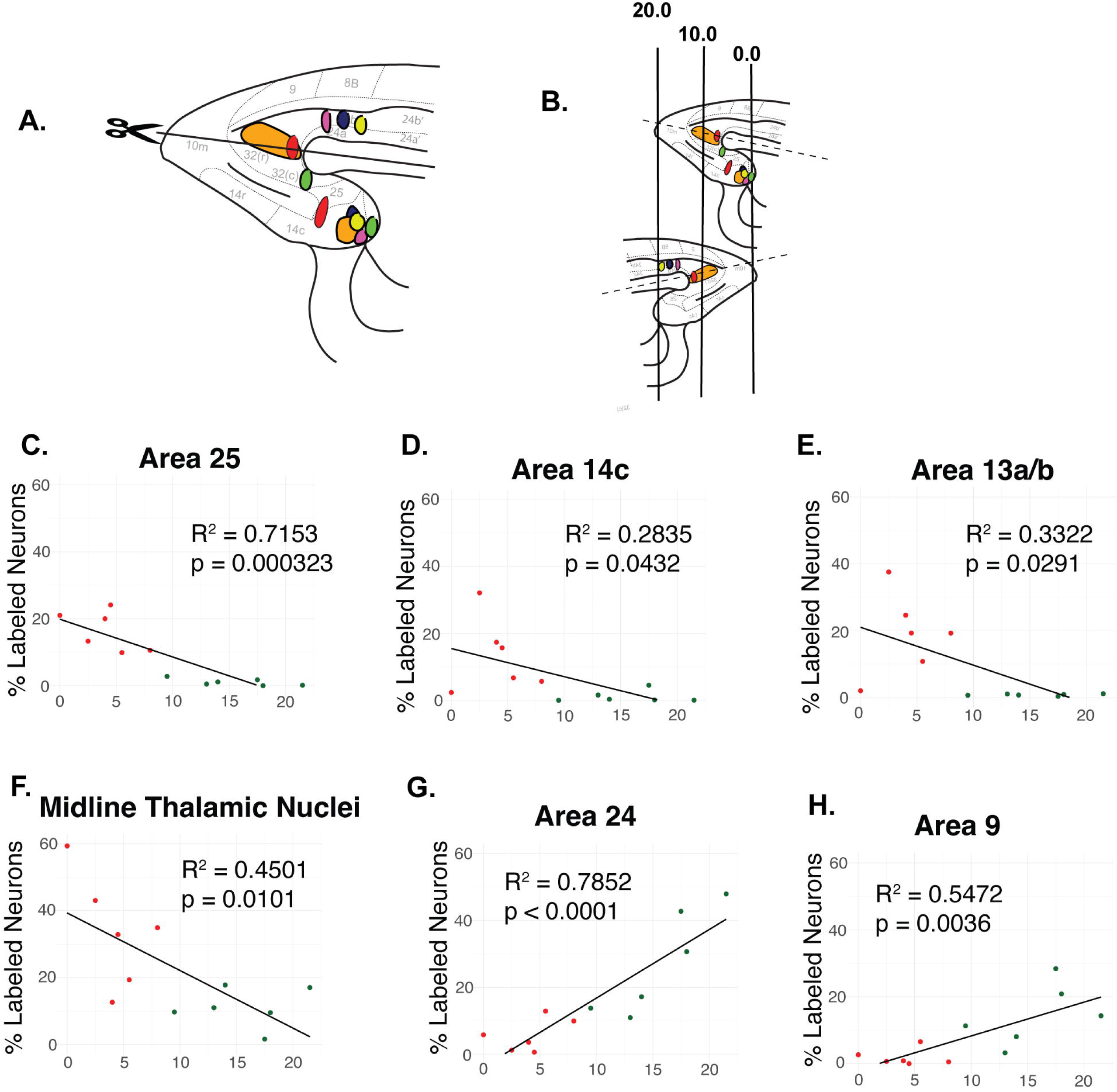
Afferent pathways that correlate with linearized anatomy of the ACC. Red = sgACC site, Green = pgACC site. A-B. Sagittal maps depicting site location in standardized space after ex vivo mapping. C. Linear regression between the linearized injection site coordinates and the percentage of total labeled PFC cells found in area 25 (R^2^ = 0.7153, p = 0.000323). C. Linear regression between the linearized injection site coordinates and the percentage of total labeled PFC cells found in area 14c (R^2^ = 0.2835, p = 0.0432). D. Linear regression between the linearized injection site coordinates and the percentage of total labeled PFC cells found in area 13a/b (R^2^ = 0.3322, p = 0.0291). E. Linear regression between the linearized injection site coordinates and the percentage of total labeled thalamic cells found in the midline nuclei (R^2^ = 0.4501, p = 0.0101). F. Linear regression between the linearized injection site coordinates and the percentage of total labeled PFC cells found in area 24 (R^2^ = 0.7825, p < 0.0001). G. Linear regression between the linearized injection site coordinates and the percentage of total labeled PFC cells found in area 9 (R^2^ = 0.5472, p = 0.0036).

### Correlated cortico-thalamic and cortico-cortical projections to the ACC

We next asked whether projection weights from specific brain regions tracked one another, i.e. increased or decreased their influence linearly. When all inputs to all ACC injection sites were considered, we found 4 positive correlations. Correlations between, area 25 and midline thalamic nuclei (r = 0.6643, p = 0.0219), and between area 14c and area 13a/b (r = 0.7972, p = 0.0029) were found, driven by injections in the sgACC (red dots) (**Fig. 6 A-B**). A positive relationship between proportions of labeled cells in area 24a/b and area 9 were strongly correlated across all sites (**Fig. 6C**)(r = 0.9301, p < 0.0001). A positive correlation between area 9 and VA thalamus proportions (r = 0.6119, p = 0.0345) was also noted (**Fig. 6D**). There were no correlations among any thalamo-thalamic pairs of inputs. For all 4 significant pairs, the relationship between each significant pair was driven by one ACC subdivision.

**Figure 6.**
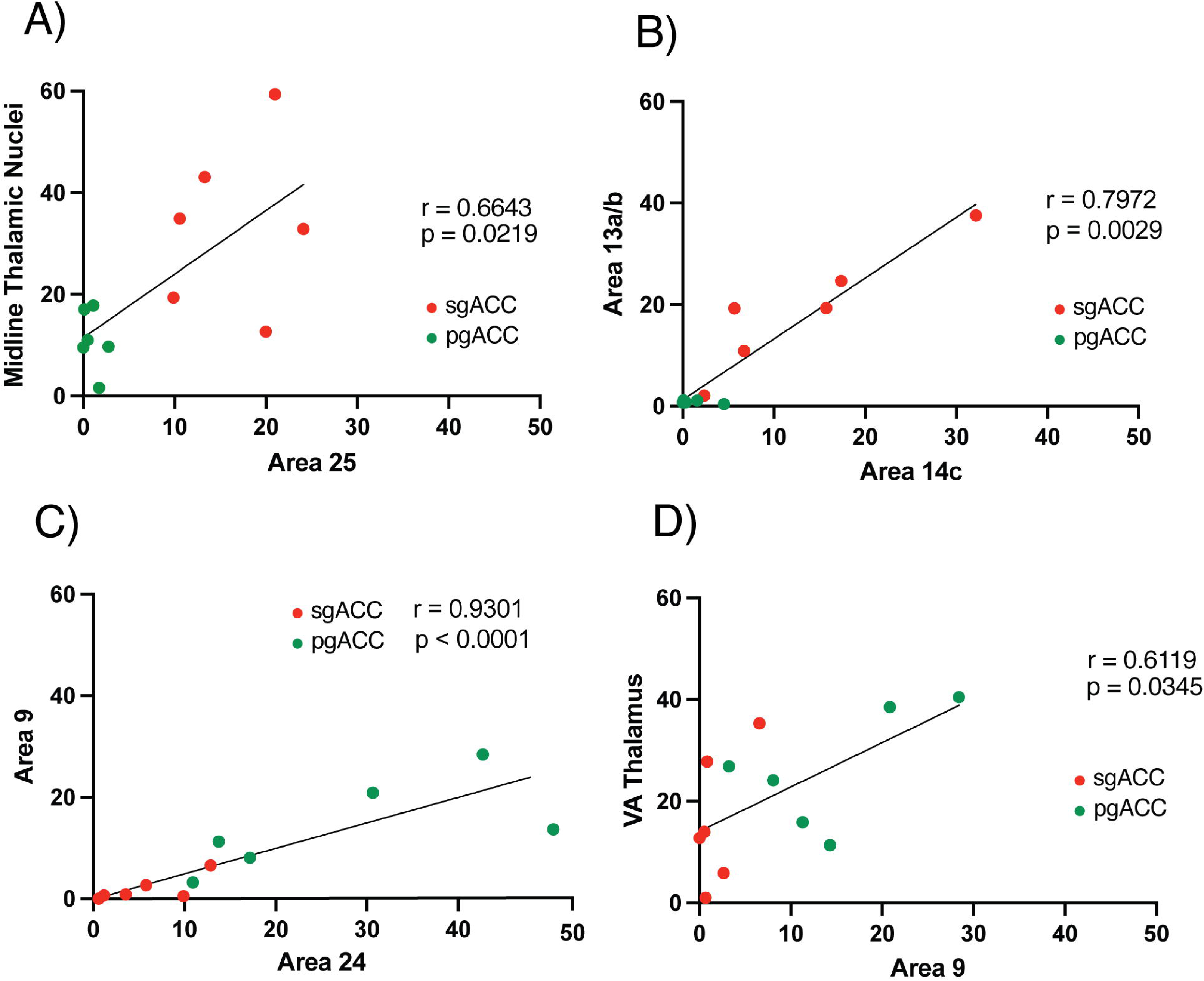
Pairs of cortical/thalamic areas project together across the ACC trajectory. A. Area 25 and the midline nuclei (r = 0.6643, p = 0.0219). B. Area 14c and 13a/b (r = 0.7972, p = 0.0029B. C. Areas 24 and area 9 (r=0.9302, p<0.0001). D. Area 9 and VA thalamus (r=0.6119, p=0.0345). Red = sgACC site, Green = pgACC site.

### Organization of cortical and thalamic afferent connectomes to sgACC and pgACC

To understand the balance of all cortical inputs to the sgACC versus pgACC sites, we assessed the proportion of labeled neurons in all cortical areas, sorted by a granularity index that we previously devised(McHale et al., 2022) **(Fig. 7).** We did the same for thalamic nuclear regions, organizing them by their connectivity with cortical regions as previously described.

**Figure 7.**
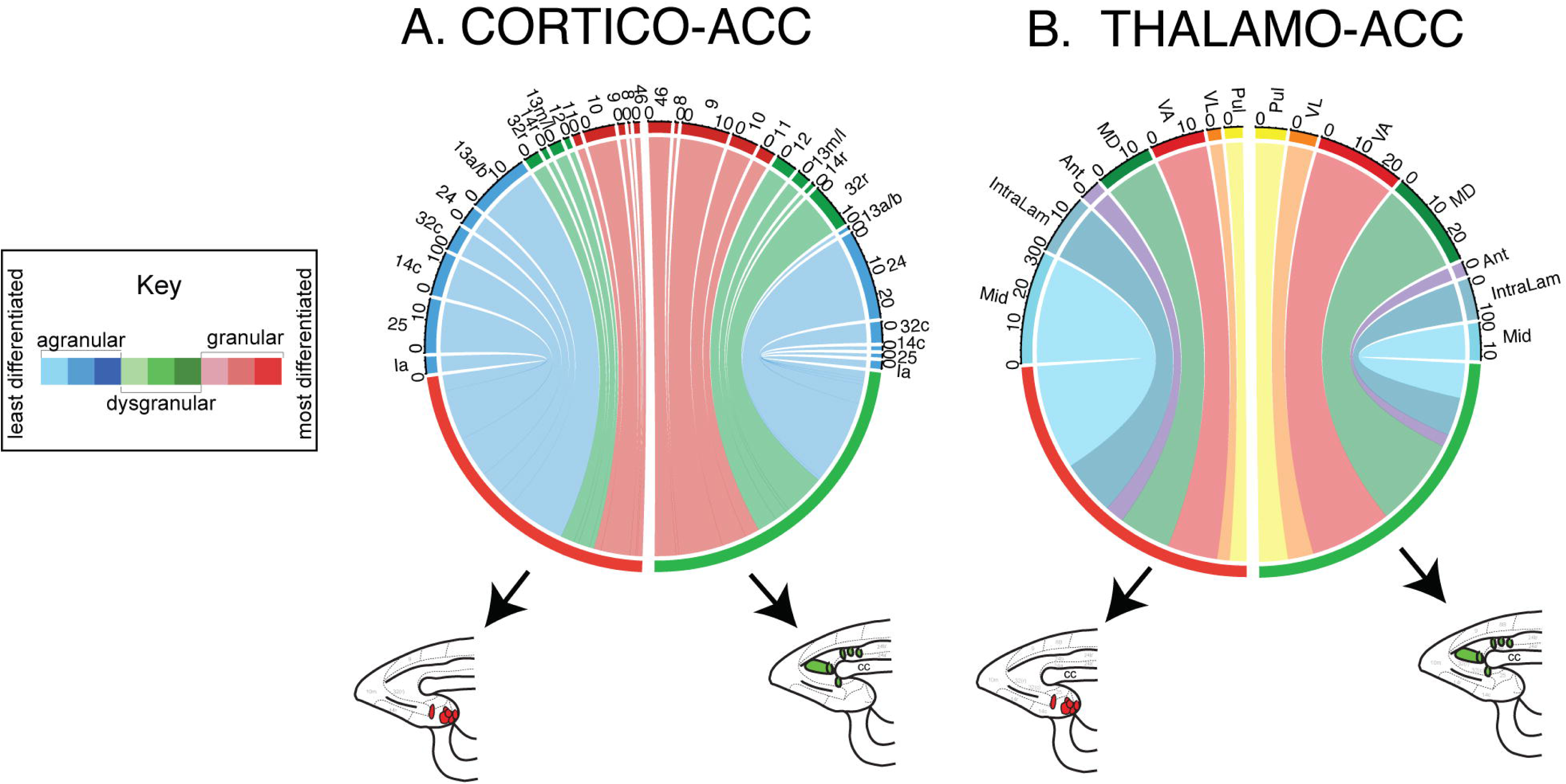
Chord plots showing the flow of projections to the sgACC and pgACC. A. Hemi-chord plots showing the average percentage of inputs from different cytoarchitectural regions after all injections to the sgACC (left) and pgACC (right). Cortical areas are color coded by granularity assignment, Blue = agranular, Green = dysgranular, Red = granular. B. Chord plot showing the average percentage of inputs from thalamic regions after all injections to the sgACC (left) and pgACC (right). Thalamic areas are color coded corresponding to the cortical areas with which they have known connections, *Light Blue = midline, Dark Blue = intralaminar, Purple = anterior, Green = mediodorsal, Red = VA, Orange = VL, Yellow = pulvinar*.

Both the sgACC and pgACC received broad inputs from agranular, dysgranular and granular cortices. However, the relative weights of these cortical regions were strikingly different in the sgACC versus the pgACC. Agranular cortical areas constituted a greater proportion of total inputs to sgACC (blue: 70%) accompanied by inputs from dysgranular (14%) and granular cortical sites (16%) (**Fig. 7A**). The proportion of afferent contributions from the agranular cortices was markedly reduced for pgACC sites (blue: 40%, or approximately 57% of the total in sgACC), and replaced by proportionate increases in dysgranular (green: increasing from 14% for sgACC to 26% for pgACC) and granular cortical areas (red: more than doubling from 16% in sgACC to 34% in pgACC (**Fig. 7A**). Overall, pgACC received a more even distribution of afferents from all laminar categories.

For the thalamic nuclear groups, we also found that both the sgACC and pgACC received an array of inputs from all thalamic regions analyzed. The relative contributions of each thalamic nuclear group, when organized by cortico-thalamic connectivity, mirrored the findings for the cortex. Thalamic nuclear areas (midline, blue; intralaminar nuclei, dark blue) known for connections to the agranular cortex (Jones, 1985; Hsu and Price, 2007) constituted 52% of all inputs to the sgACC, compared to MD (green, 16%) and VA (red, 16%) which are relatively more interconnected with dysgranular and granular regions of the PFC (Goldman-Rakic and Porrino, 1985; Jones, 1985; McFarland and Haber, 2002) (**Fig. 7B**). In the pgACC, midline and intralaminar inputs were proportionately decreased compared to sgACC sites (blue: 52% in sgACC to 22% in pgACC), and replaced with increased proportions of labeled cells in MD (green: 16% in sgACC to 28% in pgACC) and VA (red: 16% in sgACC to 26% in pgACC) (**Fig. 7B**). Thus, the relative balance of inputs to sgACC and pgACC can be predicted by cortical granularity, with the balance of thalamic inputs showing a similar pattern, derived from known cortico-thalamic connections.

## Discussion

This study takes a mesoscopic approach to examine the PFC and thalamic afferent networks of the ACC functional subdivisions. Several important findings emerge. First, the sgACC and pgACC indeed receive many common cortical and thalamic afferents, but are distinguished by the relative weights (‘drivers’) of specific cortical and thalamic regions. Unbiased clustering showed that sgACC and pgACC ‘driver’ systems were almost wholly dissociated. In general, the sgACC was under greater direct influence of areas important for encoding value and maintaining autonomic arousal, such as area 25, 14c, 32c, area 13a/b, and the midline thalamic nuclei (Rudebeck and Murray, 2011; Hsu et al., 2014; Murray et al., 2015). The pgACC received the most heavily weighted inputs from regions involved in executive function, including working memory and set-shifting, such as dorsolateral PFC (dlPFC) areas 9 and 46, and the MD and VA thalamus (Friedman and Goldman-Rakic, 1994; Petrides, 2000; Wolff and Halassa, 2024; Phillips et al., 2025). Area 10m, a region found only in primates, is the only common ‘driver’ of both the sgACC and pgACC. It has a role in ‘self-generated’ decision-making and exploration, suggesting that ‘self’ percepts influence both ACC regions (Tsujimoto et al., 2010; Mansouri et al., 2017).

Second, ‘modulatory’ circuitry emerged as an important source of variability with respect to individual injection sites. Interestingly, some but not all variability was due to caudal to dorsal placement of injections in the linearized ACC. Areas 25, 14c, 13a/b, 24a/b, 9 and the midline thalamus showed a significant stepwise change in projection weights moving over the ACC trajectory. We interpret small changes in these projection weights to be related to incremental changes in ACC cytoarchitecture. The lack of linearity correlations in other projections may suggest a more general and/or spotty projection to the entire region (e.g. the pulvinar)(Romanski et al., 1997).

Finally, when examined together, afferent connectomes to the ACC were dictated by an organizing principle based on laminar differences both at the target (sgACC versus pgACC) and from afferent sources (**Fig. 8**). The agranular sgACC has a different balance of inputs than the slightly more differentiated pgACC. These principles extend to thalamic inputs, which follow the pattern of their main cortical partners, underscoring the important dependency of the cortex on the thalamus and vice versa (Anton-Bolanos et al., 2018; Steiner et al., 2020).

**Figure 8.**
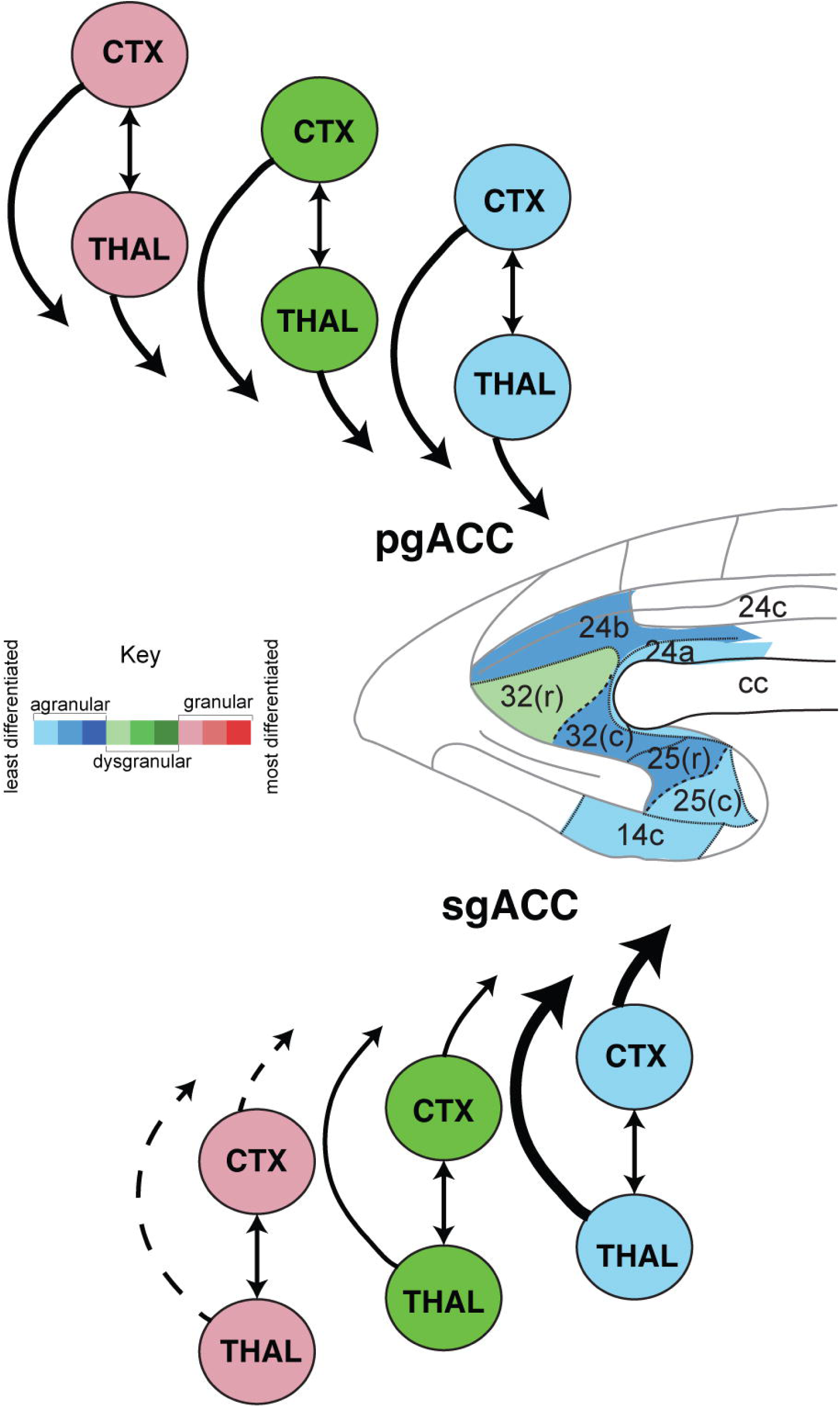
Cortical and thalamic interconnectivity links afferent weighting of both projections to both the sgACC and pgACC.

### Individual tracing studies

Our results replicate and expand on the findings of many previous circuit tracing studies in single animals. For example, previous work identified dense retrograde labeling in the orbital cortex after retrograde tracer injection into the sgACC (Carmichael and Price, 1996; Joyce and Barbas, 2018), which were recapitulated, with the additional finding that orbital areas 13a and 13b are “drivers” of the sgACC, but not the pgACC. Orbital cortical ‘drivers’ to the sgACC may be essential for encoding of reward value, given that lesions selectively disrupt encoding of object value on a series of reward-related tasks (Rudebeck and Murray, 2011). We also identified dlPFC areas 9 and 46 as “drivers” of the pgACC, but not sgACC. These results are consistent with previous individual circuit tracing studies (Barbas and Pandya, 1989; Carmichael and Price, 1996). Together, these findings have long supported pgACC’s role in higher-order cognition, like decision-making and conflict monitoring (Modirrousta and Fellows, 2008; Livneh et al., 2012; Apps et al., 2016; Dal Monte et al., 2020).

We found that frontopolar area 10m was a common “driver” of both sgACC and pgACC. This result is also consistent with earlier anterograde and retrograde studies in macaque (Carmichael and Price, 1996; Ongur and Price, 2000; Petrides and Pandya, 2007), and with a probabilistic tractography study in human (Liu et al., 2013). Our connectomic approach extends these findings by showing that area 10m stands out as the sole ‘driver’ for both sgACC and pgACC nodes, while other ‘driver’ systems are largely dissociated. This dual ‘driver’ status places area 10m as an important portal for ‘top-down’ information flow to both ACC subregions. Area 10m integrates value representations with self-generated goals, and may be an important afferent during tasks that require emotional valuation and introspective control of cognition (Hogeveen et al., 2022).

Our results are also in agreement with individual thalamic circuit tracing studies. It is well known that the midline thalamic nuclei are reciprocally connected with the sgACC (Barbas et al., 1991; Hsu and Price, 2007). We found similar results, and noted that the midline thalamic nuclei inputs is strongest caudally, tapering rostro-dorsally. This connection provides arousal information based on relays from autonomic brainstem regions such as the periaqueductal gray, the parabrachial nucleus, and the locus coeruleus (Penzo and Gao, 2021). Minor inputs to the sgACC from VA and MD is also consistent with previous work (Akert and Hartmann-von Monakow, 1980; Vogt et al., 1987; Barbas et al., 1991; McFarland and Haber, 2002). The pgACC was mainly innervated by the MD and VA thalamus, which is in line with previous studies (Giguere and Goldman-Rakic, 1988; Barbas et al., 1991; McFarland and Haber, 2002). The VA and MD thalamus work sequentially through the cortex to support higher cognition, based on recent physiologic studies in macaque (Phillips et al., 2025). VA first encodes and transmits an abstract rule to the cortex, whose representation is maintained through reciprocal MD-PFC loops that map onto motor selection (Phillips et al., 2025). Strong inputs from VA and MD to pgACC may suggest similar contributions to hierarchical decision-making, including in social contexts (Amemori and Graybiel, 2012; Chang et al., 2013; Apps et al., 2016).

### Cortex and thalamus connectomes project in tandem

Our previous work demonstrated that cortical laminar differentiation dictates a hierarchical set of inputs in PFC-amygdala and PFC-striatal circuits (Cho et al., 2013; McHale et al., 2022). In these studies, the agranular cortex formed a broad base of afferent terminals, with progressively more differentiated cortical regions terminating in step-wise manner along a topographic gradient. Thus, agranular cortical terminations co-projected with dysgranular cortex, followed by the addition of granular cortical inputs in adjacent targets. Applying that same analysis here, we found that the rich variety of inputs from agranular, dysgranular, and granular cortex followed similar patterns. The slightly less differentiated sgACC receives a greater share of agranular PFC inputs than pgACC. In contrast, pgACC (areas 24/32), which has a relatively more organized cytoarchitecture (including incipient layer IV in some parts), had a shift, receiving agranular inputs, but with a greater portion of dysgranular and granular PFC inputs than sgACC. These results provide insight into how cortical information is combined in each region. The thalamus is essential to cortical function, forming a link to basal ganglia output circuitry. When organized according to its main cortical partners, the thalamus created similar maps, highlighting the tight relationship between these two afferent systems.

### ’Top-down’ relationships dominate the ACC: clinical applications

One question was whether sgACC and pgACC are intrinsically connected. Surprisingly, we found that area 32 was a ‘driver’ of the sgACC, but neither areas 25 or 14c were drivers of the pgACC. There is thus a pgACC “top-down” input to the sgACC, but no direct “bottom-up” input to pgACC from the sgACC. One study in rodents using tracing and microstimulation reports similar results (Marek et al., 2018). This flow of intrinsic connectivity comported with the overall flow of cortical connections to both the sgACC and pgACC, which was generally incremental, from next level differentiated cortices down to their respective targets (**Fig. 3**). These observations may be useful for considering strategies to optimize neuromodulation therapies for MDD. Transcranial magnetic stimulation (TMS) of the dlPFC has antidepressant effects, and its clinical efficacy is thought to be related to alterations in sgACC activity (Pascual-Leone et al., 1996; Fox et al., 2014; Liston et al., 2014). However, how dlPFC regional stimulation drives sgACC engagement is unknown, since monosynaptic connections between the dlPFC and sgACC do not exist (Joyce and Barbas, 2018; Joyce et al., 2020; Cash and Zalesky, 2024). Therefore, dlPFC stimulation likely propagates to the sgACC polysynaptically. Our work supports that dlPFC areas 9 and 46 serve as driving afferent inputs of the pgACC, with pgACC area 32 directly projecting to the sgACC. Another possibility for dlPFC engagement includes the known connection to the area 10m (Barbas and Pandya, 1989; Rosa et al., 2019), which is a ‘driver’ of both ACC subdivisions based on our results. Together our findings suggest testable routes of neurotherapeutic engagement of the sgACC and pgACC.

## Conclusion

In conclusion, these results demonstrate that defining afferents of ACC subdivisions are related to a set of hierarchical rules, which are governed by cortical granularity and thalamo-cortical projections. The ACC subdivisions are themselves heterogenous cortical regions with afferents “added” and “subtracted” as their cytoarchitecture itself becomes more differentiated along the ACC trajectory. These findings may not only have implications for understanding the connectional basis of sgACC and pgACC function, but also may be relevant for understanding the neural basis of psychiatric disorders and improving targeting for neuromodulation therapies for MDD.

## Acknowledgements

Supported by R01MH130608 (JF), T32NS115705 (DM). This work was also supported by Center for Advanced Brain Imaging and Neurophysiology at the URMC.

## Author Contributions

DCM performed research, analyzed data, and wrote the first draft of the manuscript. TML aided in statistical analyses. JLF designed the research, supervised data collection and analysis, and edited the manuscript.

